# Early identification of cooperative fragments for protein-protein interaction stabilization

**DOI:** 10.1101/2022.02.23.481635

**Authors:** Pim J. de Vink, Peter J. Cossar, Bente A. Somsen, Christian Ottmann, Luc Brunsveld

**Affiliations:** Laboratory of Chemical Biology, Institute for Complex Molecular Systems and Department of Biomedical Engineering, Eindhoven University of Technology, P.O. Box 513, Eindhoven, 5600 MB, The Netherlands

**Keywords:** Protein-Protein Interactions, Fragment-based drug discovery, Saturation transfer difference NMR, cooperativity, molecular glues, 14-3-3

## Abstract

Modulating protein-protein interactions (PPIs) is an effective approach to drug discovery, with several drugs in the clinic that inhibit PPIs. The orthogonal approach of PPI stabilization has developed slowly, a function of the complicated dynamics of multi-component protein complexes. In contrast to PPI inhibition, where ligand affinity is the driving parameter for efficacy, cooperativity is frequently the directing variable for PPI stabilization. Here we show how STD NMR allows for early-stage detection of cooperativity using the hub protein 14-3-3, a focused library of fragments and several 14-3-3 partner proteins. Further, we validate that the observed enhancement in STD signal is a function of cooperativity of the ternary 14-3-3 complex, using mutagenesis and X-ray crystallography. Additionally, we assess the differential cooperativity of three fragments in a panel of 14-3-3 interaction partners. Finally, we demonstrate how selective 14-3-3 complex formation is a function of cooperativity effects

## Manuscript

Stabilization of protein-protein interactions (PPIs) with small molecules is an emergent strategy in drug discovery with the potential to target 'hard to drug' protein classes^1^. PPI stabilizers, such as molecular glues, enhance the affinity of a protein complex through a cooperativity mechanism. Cooperativity is a common phenomenon in complex biological systems, where an initial binding event enhances the affinity of subsequent binding events^2,3^. Both synthetic PPI stabilizers, like Pleconaril and Tafamidis, as well as natural products, have shown remarkable target selectivity, with Rapamycin and Taxol used in the clinic for decades. Despite these successes, molecular glues remain challenging to design *de novo*, with the vast majority discovered retrospectively^4^. Notable exceptions are bifunctional compounds such as PROteolysis-TArgeting Chimeras (PROTACs)^5,6^. This approach to targeted protein degradation is partly converging with PPI stabilization towards more cooperative compounds^7^.

A significant challenge for the (early stage) discovery of new cooperative compounds is the complicated dynamics of multi-component complexes of proteins and small molecules. Not only is the affinity of a compound important, but cooperativity is similarly critical to protein complex stabilization. This requires hit compounds in a screening campaign to pass through a double selection filter of occupancy (affinity) and ability to induce stabilization (cooperativity) to elicit an assay response. This combination of criteria is particularly challenging to meet for screening libraries, given the weak affinities of initial PPI stabilizer hits, and is further amplified for fragment screens^8^.

To circumvent the challenges of identifying low affinity chemical matter, covalent fragment approaches have been developed^9^. Covalent tethering typically employs a reactive group that anchors a fragment on a protein surface^10^. Our laboratory and others have further developed this approach for PPI stabilization. Specifically focusing on reversible covalent tethering approaches that exploit disulfide^11^ and imine chemistry^12^ to anchor the fragments at the PPI interface. Notwithstanding, removal or substitution of the reactive group remains challenging due to molecular recognition of the fragment being dependent on the reactive group or the specific molecular orientation induced by the covalent linker^13^. Therefore, non-covalent fragment libraries also have significant value, given hit detection is dominated by molecular recognition^14^.

Fragment-based drug discovery has proven to produce clinical candidates via fragment growth, linking, merging or self-assembly strategies using non-covalent fragments as initial starting points^15^; and provides a solid foundation of medicinal chemistry principles to improve ligand potency^16,17^. In contrast, our understanding of optimizing cooperativity is still developing^18,19^. Therefore, biophysical approaches to detect cooperativity at the early stages of hit identification are urgently needed.

Most standard biophysical screening techniques, such as Fluorescence Anisotropy (FA), Förster Resonance Energy Transfer, Isothermal Calorimetry, and Surface Plasmon Resonance, are optimized to detect relatively strong binding interactions. The affinity detection windows of these techniques are less suited for initial PPI stabilizer hit identification, especially of the fragment-type^20,21^. Further, many fluorescent biophysical assays have an indirect output which can decrease the observed cooperativity in these assays^2^. Also, certain fragments (termed *silent binders*) can form ternary complexes in X-ray crystallography, while having no effect in solution-based assays^12^. There is a significant need to develop assays to identify cooperativity that function at the low millimolar range to detect early PPI stabilizer chemical matter.

Saturation Transfer Difference (STD) NMR, waterLOGSY^22^ and ^19^F NMR can be used to detect millimolar potent fragments^23,24^. STD NMR has been used for example successfully to observe allosteric cooperativity for GPCRs^25,26^. Furthermore, STD NMR is ligand-observed and measures the binding event of the ligand directly. Also, within the STD NMR assay format PPI partners can be combined at concentrations that allow for high binary PPI occupancy. NMR is thus well suited to measure the weak affinities associated with fragments^27^, monitoring ligand binding from the preformed binary PPI complex.

Utilizing 14-3-3 and a panel of its phosphorylated partner peptides as a case study, we characterize early stage cooperativity in fragment binding using STD NMR. 14-3-3 proteins are involved in numerous cellular functions, including scaffolding, nuclear occlusion and signal transduction^28^. The protein typically binds disordered regions of phosphorylated partners via conserved phorphorylated serine or thrionine epitopes^29^. Given the extensive 14-3-3 proteome, consisting of hundreds of binding partners, stabilization of a specific PPI is crucial for therapeutic benefits. We demonstrate that 14-3-3 complex stabilization can be detected at low mM concentration using STD NMR. We also show that differential cooperativity influences the selectivity of complex formation, leading to early-stage detection of selective complex formation.

Previously, using X-ray crystallography, we identified that amidine substituted thiophenes and benzothiophenes bind into the phosphopeptide-binding-groove of 14-3-3, adjacent to the TAZ phosphopeptide (Fig. 1a)^30^. The amidine moiety of **AZ130**, which is essential for activity in this class of compounds, forms a salt bridge with E14 of 14-3-3σ. However, **AZ130** has no enhancing effect on the 14-3-3:TAZ interaction in FA assays. Utilizing **AZ130** as a model fragment, we sought to investigate the binding of **AZ130** with and without the TAZ peptide to assess the capacity of STD NMR to detect cooperative fragment binding (Fig. 1b). In STD NMR, the receptor protein and peptide are selectively presaturated to suppress their signals.^23^ This pre-saturation is transferred to binding fragments resulting in a decrease of the signals associated with the binding fragment. We propose that synergistic effects of the peptide in the presence of 14-3-3 and the fragment will translate to a more pronounced STD signal for the ligand. Therefore, an STD experiment was performed in the presence and absence of the partner peptide TAZ (10-fold excess). Important to us was the tractability of this methodology for other research groups. We were particularly interested in applying this technique to a conventional 400 MHz NMR machine without the need for a cryoprobe. Screening of **AZ130** was conducted on a Bruker 400 MHz NMR with *apo*-14-3-3σ and the 14-3-3σ/TAZ complex. Each STD experiment was conducted at 300 *μ*M of **AZ130**, 10 *μ*M of 14-3-3σ in PBS buffer and 10% D_2_O. Analysis of the STD spectrum of the **AZ130** with 14-3-3σ identified two weak proton signals at 7.35 and 7.65 ppm, consistent with the protons associated with the phenyl and thiophene rings of **AZ130**, suggesting that **AZ130** shows weak binding to 14-3-3σ alone (Fig. 1c). Analysis of the 14-3-3σ/TAZ complex featuring 100 *μ*M TAZ peptide and **AZ130** (300 *μ*M) revealed significantly enhanced the proton signals, implying cooperative complex formation. This result was consistent with the observed X-ray crystal structure of **AZ130** in complex with 14-3-3/TAZ.

**Figure 1.**
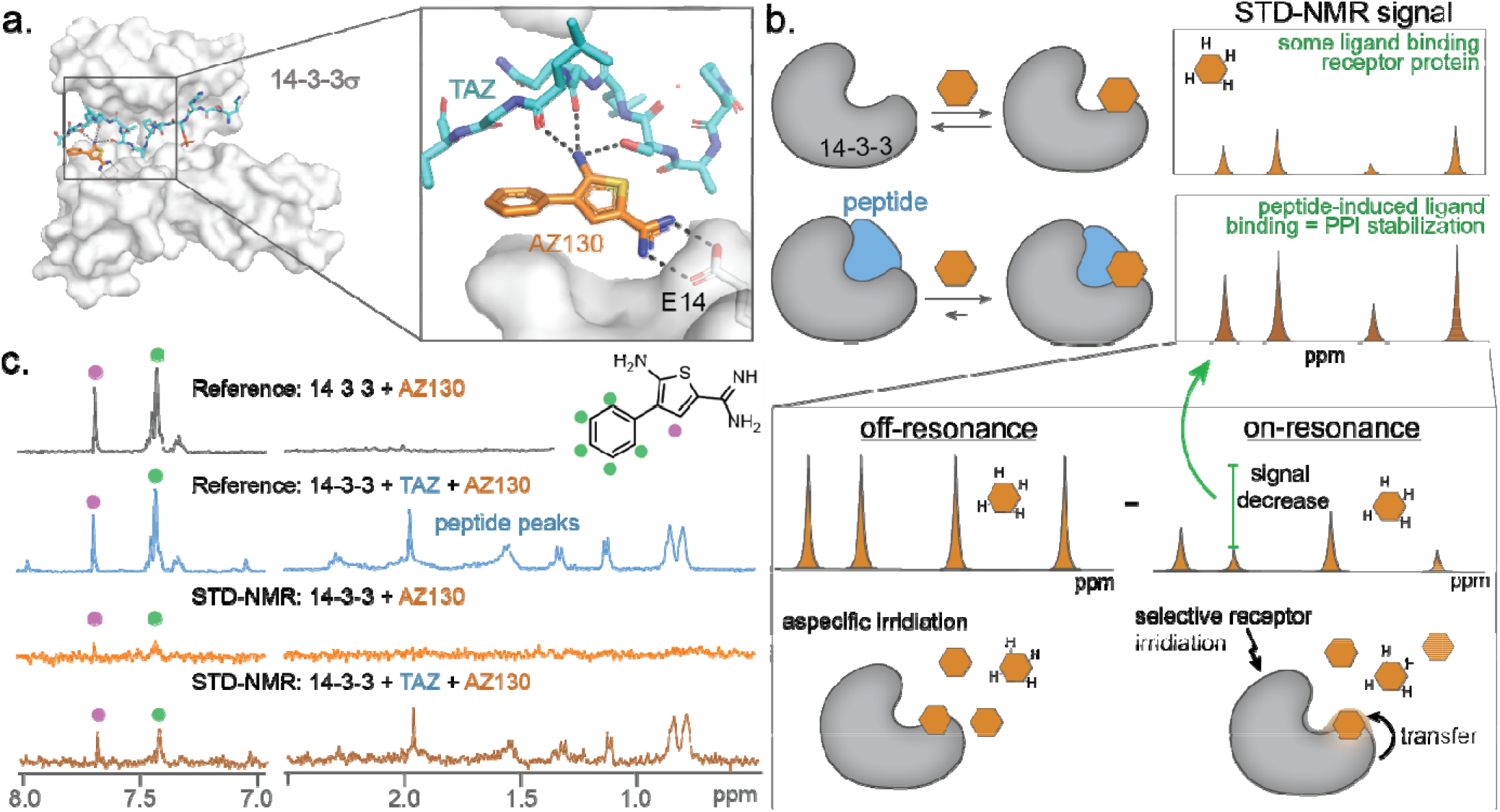
Detection of cooperative fragments by STD NMR. **a.** X-ray crystal structure of **AZ130** in complex with 14-3-3σ/TAZ peptide (PDB:6RHC). An enlargement of the 14-3-3/TAZ/AZ130 interface is shown in the grey box. 14-3-3 (grey) is depicted with Van Der Waals surface; TAZ peptide (cyan) is shown as a stick model; **AZ130** (orange) is shown as a stick representation; polar interactions are depicted as black dashed lines. **b.** Schematic representation of enhanced STD signal due to ternary complex formation between the 14-3-3 (grey), partner peptide (blue) and fragment (orange). The bottom panel depicts a cartoon representation of how an STD NMR spectrum (off-resonance – on-resonance spectrums) is derived, which is more pronounced in the case of the ternary complex. **c.** Reference and STD NMR spectra of 300 *μ*M **AZ130** with 10 *μ*M 14-3-3σ in the absence or presence of 100 *μ*M TAZ-peptide in PBS (50 mM NaH_2_PO_4_/Na_2_HPO_4_ pH 7.5, 150mM NaCl, 10% D_2_O).

Having confirmed the application of STD NMR to detect low affinity fragment binding to the 14-3-3σ/TAZ complex, as well as indications of cooperativity, we investigated a focused library of amidine derivatives (Fig. 2a). The amidine functional group was maintained, given its important electrostatic interaction with 14-3-3 residue E14. Notably, this library elicited no response in a FA assay (Supporting Fig. S1). The focused library was screened using STD NMR against the *apo*-14-3-3σ protein and the 14-3-3σ/TAZ complex. Analysis of the nine fragments identified that **F1, F2, F4, F5, F8** and **F9** all elicited assay responses. Of these fragments, five bound to *apo*-14-3-3 (Fig. 2b, Supporting Table S1). Fragments **F2, F4, F5** and **F9** showed enhanced signals in the presence of the TAZ peptide (see supplementary NMR spectra). **F2, F4** and **F5** elicited an improved binding of 5.3−, 1.9− and 2.2-fold, respectively, in the presence of TAZ (Fig. 2c & d). Notably, fragment **F9** did not bind to *apo*-14-3-3; however, a significant signal was observed upon the addition of the TAZ peptide. The fragments **F3, F6** and **F7**, showed no signals for their aromatic protons in either the *apo*-14-3-3σ or 14-3-3/TAZ complex STD spectra, indicating no fragment binding in either case. A decrease in signal for **F1** and **F8** was observed in the presence of TAZ, indicating competition for 14-3-3 biding with TAZ for these fragments (Fig. 2b, c & e).

**Figure 2.**
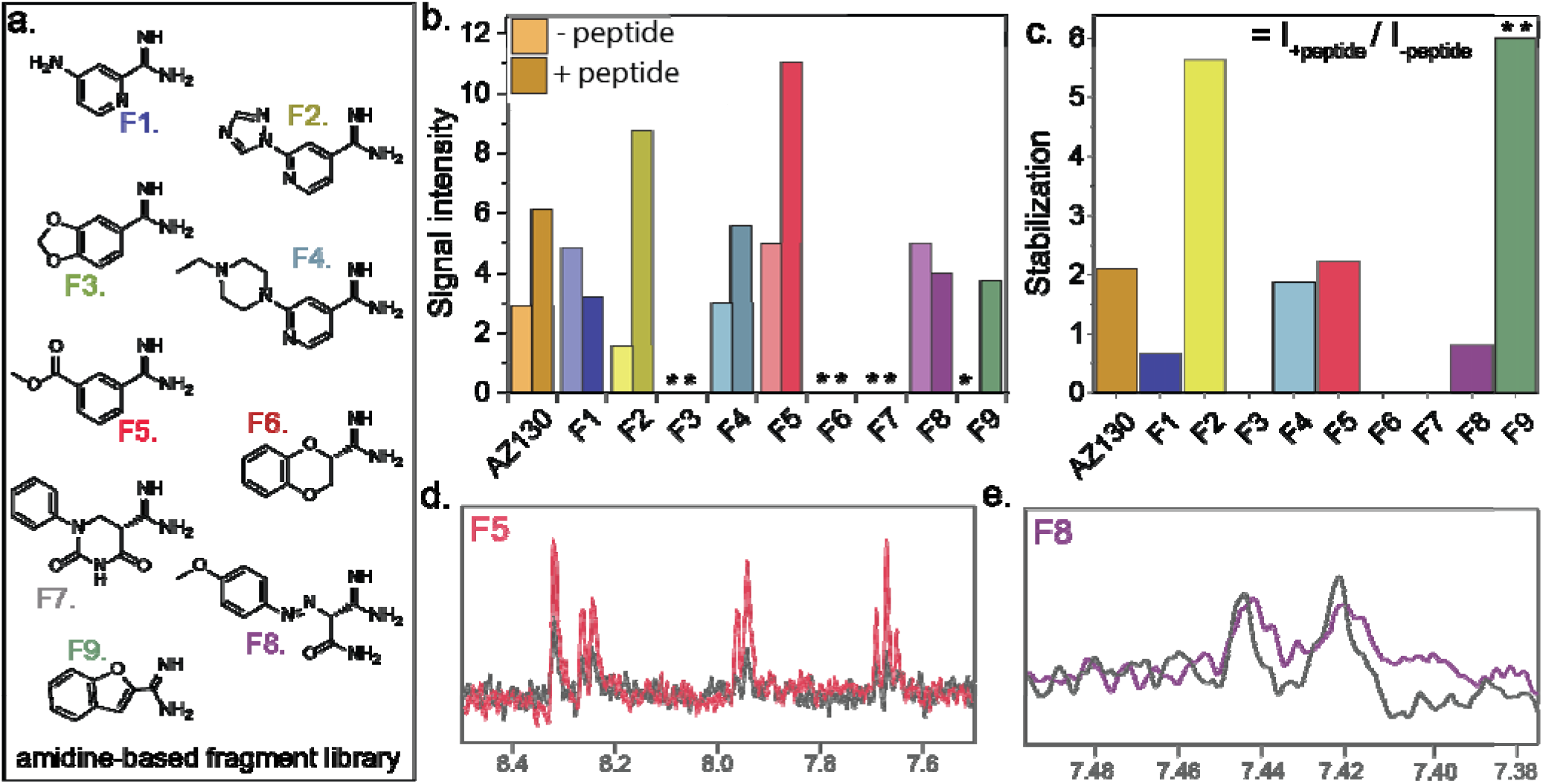
Initial fragment screening against TAZ. **a.** Focused library of small molecule amidine analogues. **b.** STD signal of fragments (300 *μ*M) with 14-3-3σ (10 *μ*M) in the absence and presence of 100 *μ*M TAZ peptide. Signal intensity was calculated based on the peak intensity of an aromatic proton relative to the baseline of the STD NMR. * indicates no signal detected. c. Fold stabilization in signal upon addition of the peptide relative the binary experiment (I_+peptide_ / I_−peptide_) ** indicates stabilization cannot be well quantified, given the fragment does not bind to *apo*-14-3-3. **d.** Comparison of raw STD signal for **F5** in the absence (black) and presence (red) of the TAZ binding partner. **e.** Comparison of raw STD signal for **F8** in the absence (black) and presence (purple) of the TAZ binding partner.

We repeated the screen of **F4, F5** and **F9** at a higher concentration (5 mM) to validate the initial screening result, using an amplification factor (A_STD_) as a concentration independent metric. The STD amplification factor was derived by multiplying the relative STD effect (*I*_on-resonance_ / *I*_off-resonance_) at a fixed fragment concentration where the molar ratio of the fragment is in excess relative to the protein (see supplementary equation 1). Analysis of fragments **F4, F5** and **F9** with wild-type 14-3-3σ was consistent with previous observations (Fig. 3a). In contrast, 14-3-3σ E14A, a mutant without the crucial glutamic acid, elicited no significant A_STD_ signal when treated with fragments **F4, F5** and **F9**, neither in the presence nor absence of the TAZ peptide.

**Figure 3.**
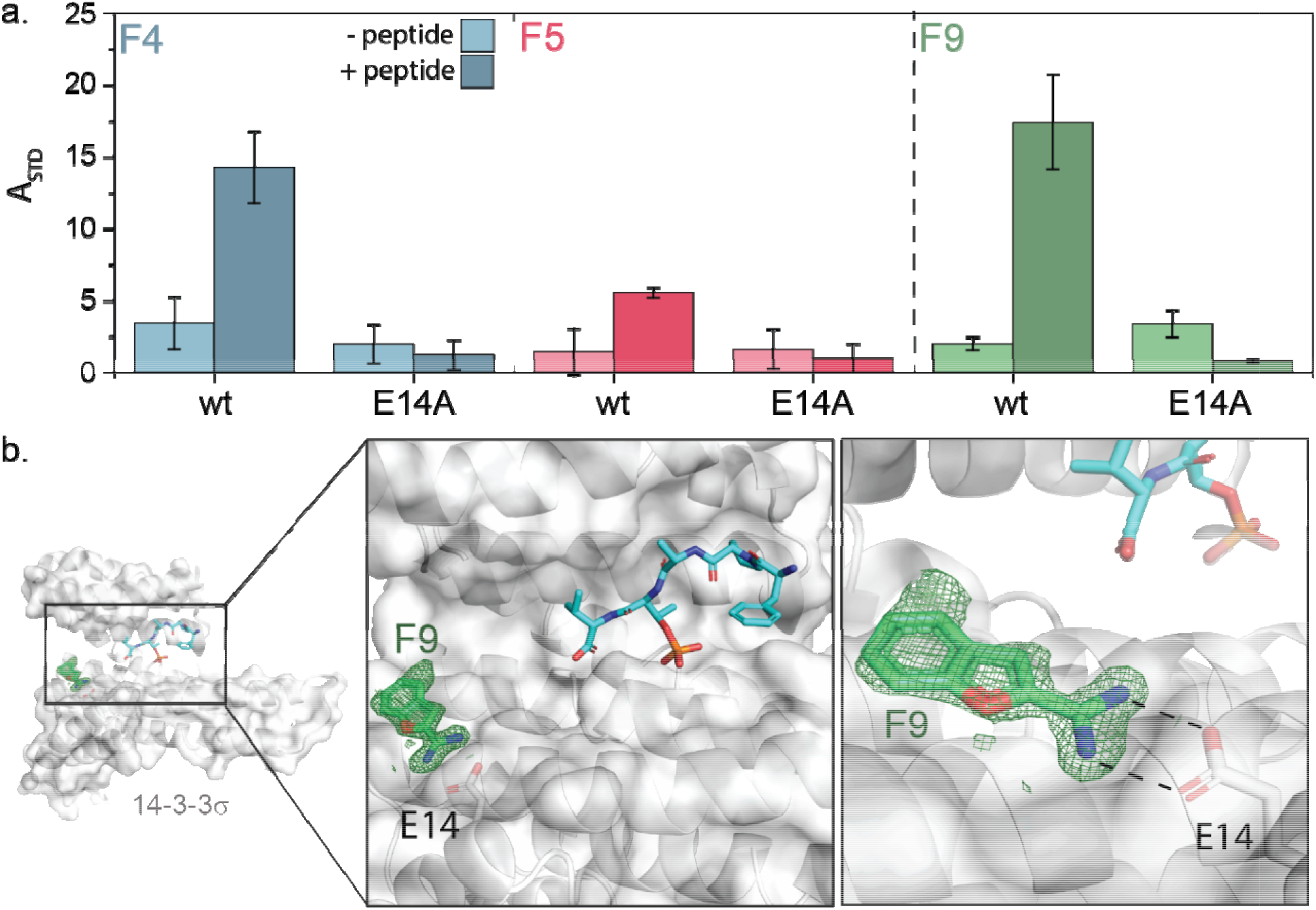
Validation of identified hit fragments. a. Comparison of enhancement of ASTD signals obtained for F4, F5 and F9 with and without TAZ peptide. For each fragment the activity using wildtype 14-3-3σ and the E14A mutant 14-3-311⍰ are compared. Conditions: 5 mM fragment, 100 μM TAZ, 10 μM 14-3-3. Error bars indicate the mean ± SD from protons involved. b. Ternary X-Ray crystal structure of F9 in complex with 14-3-3σ and an ERα peptide. Within the black box is shown a close-up of the 14-3-3/ERα/F9 interface. 14-3-3 (grey) is depicted with Van Der Waals surface; ERα peptide (cyan) is shown as a stick model; F9 (green) is shown as a stick representation; polar interactions are depicted as black dashed lines. 2Fo-Fc electron density maps are contoured at 1⍰. PDB: 7O2J

Encouraged by these results, we soaked fragments **F4, F5** and **F9** (at 10 mM) in preformed TAZ/14-3-3σ crystals. However, we were unable to obtain a structure, likely due to the weak binding affinity requiring high concentrations of the fragment in DMSO. To further validate these results, we soaked the fragments at 10, 20 and 50 mM concentrations into binary crystals of 14-3-3/ERα complex, which tolerate higher concentrations of DMSO^31^. Soaking with 50 mM **F9** resulted in a ternary **F9**/14-3-3/ERα crystal structure at 1.5Å resolution. Analysis of this co-crystal further confirmed the binding of **F9** at the aforementioned binding site (Fig. 3b, Supporting Fig. S2).

We next assessed the differential cooperativity of fragments **F4, F5**, and **F9** with a panel of other 14-3-3 binding partners. The selected partners encompass structural diversity with some epitopes occupying the complete binding groove and others filling only a part of the binding groove. Fragments **F4, F5**, and **F9** were screened at 5 mM with 100 *μ*M peptide (Fig. 4a). Fragment **F4** results in a high A_STD_ signal with most peptide/14-3-3 combinations and appears to be quite promiscuous. In contrast, fragments **F5** and **F9** have a more distinct selectivity profile. **F5** showed a preference for ABL1 and RED binding partners. Whilst, **F9** elicited a preferred response to TASK3, ABL1 and AS160 (Fig. 4b).

**Figure 4.**
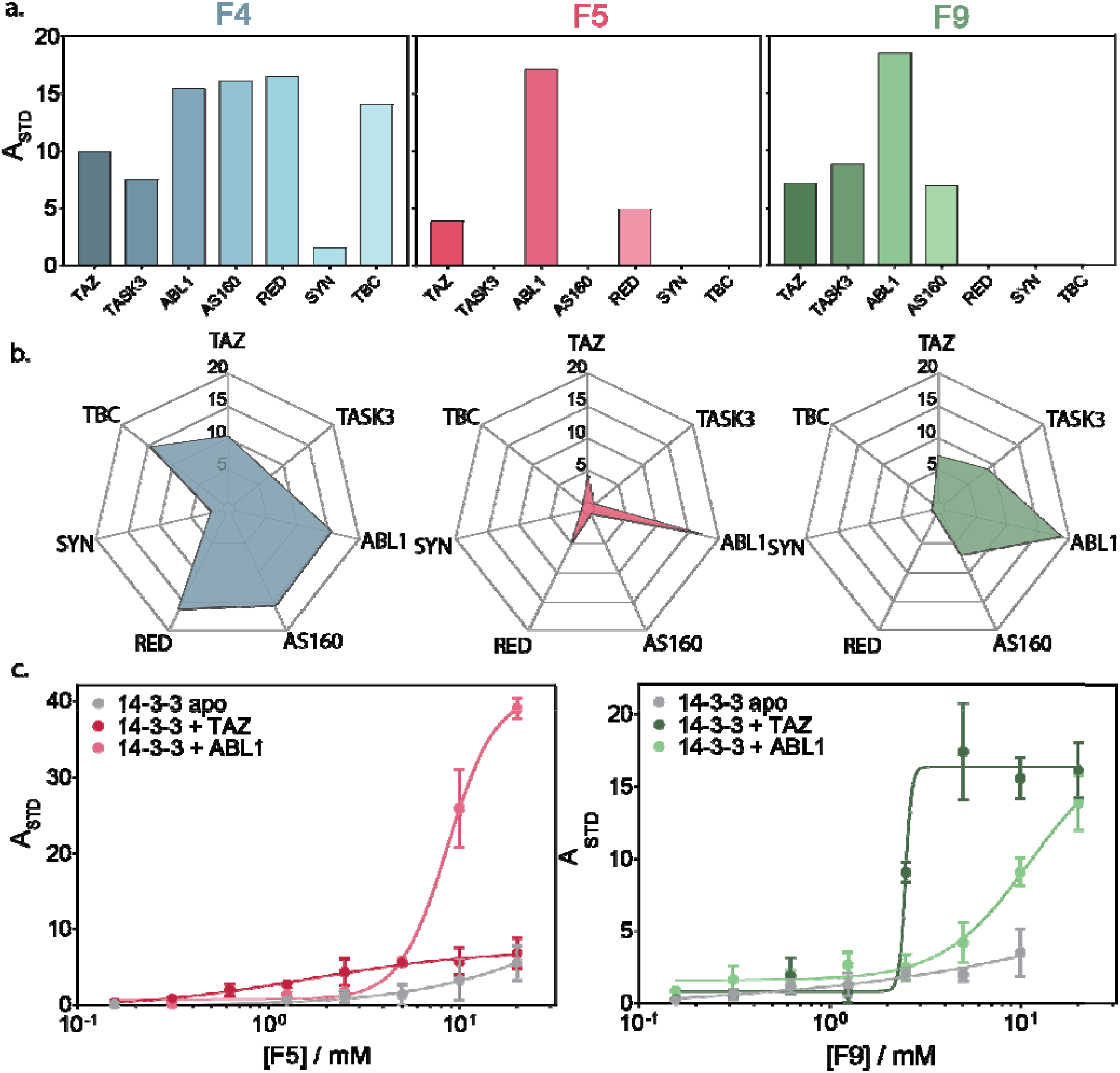
14-3-3 binding epitope selectivity screening panel. **a.** Bar chart representation of A_STD_ signal for **F4, F5** and **F9** in co-presence of various 14-3-3 binding peptides. Conditions: 5 mM fragment, 100 *μ*M peptide, 10 *μ*M 14-3-3σ. **b.** Radar plot representing the selectivity profile of each fragment. **c.** Dose response of fragments **F5** & **F9** to apo 14-3-3σ and 14-3-3σ in the presence of TAZ or ABL1. Error bars indicate the mean ± SD from protons involved.

The selected 14-3-3 binding partners TAZ and ABL1 were used to profile the dose-dependency of fragments **F5** and **F9** (Fig. 4c). The fragments were titrated to a fixed concentration of 14-3-3σ in the absence or presence of TAZ or ABL1 peptide. In the absence of a peptide only a weak signal increase was observed at higher concentrations, consistent with the fragments having very low affinity to the *apo*-14-3-3 alone. In contrast, titration of the fragments to a complex of 14-3-3/TAZ or 14-3-3/ABL1 showed a clear dose-response. Fragment **F5** elicited a dose-dependent increase in signal with EC_50_ values of 1.8 ± 0.4 and 8.7 ± 0.4 mM for 14-3-3/TAZ and 14-3-3/ABL1 complexes, respectively. Notably, variations in A_STD_ signal intensity are a function of the average number of ligand molecules saturated per 14-3-3 protein. **F9** elicited a dose-dependent increase in signal for the 14-3-3/TAZ complex with an EC_50_ value of 2.3 ± 0.3 mM. Treatment of the 14-3-3/ABL1 complex with **F9** resulted in a significant increase in A_STD_ signal. However, no saturation in A_STD_ signal was observed. Combined, these results suggest that the unique topology and functionality of the composite 14-3-3/partner protein complex drives selectivity through a cooperative mechanism with the ligand binding event.

In summary, we have shown that STD NMR is a valuable screening tool for probing early-stage cooperativity for low affinity fragments, not possible with other biophysical assays. Utilizing this approach, a modest cooperativity effect of the previously termed silent binding **AZ130** fragment was determined at high 14-3-3 and TAZ peptide concentrations. This perspective dichotomy between protein and ligand observed assays aligns with thermodynamic models for ternary systems^2^. This study further illustrates how an early focus on cooperativity in the development process can identify selective early stage hit compounds. Further, differential stabilization profiles were observed within our selectivity panel, even for these small fragments. This demonstrates that small structural modifications can significantly affect cooperativity for specific PPI combinations, forming an intriguing entry toward selectivity^32^. STD-NMR thus presents a powerful and easily accessible tool for the development of molecular glues. It enables direct measurement of fragment signals in the presence and absence of partner proteins. Further, the ubiquitous nature of NMR instruments, coupled with the technique being not dependent on isotopic enrichment, makes the approach highly amendable to both academic and industrial molecular glue research.

## Acknowledgements

This research was funded by the Netherlands Organization for Scientific Research (NWO) through Gravity program (024.001.035), Echo grant (711.017.014) and VICI grant (016.150.366) and by the European Union through the Eurotech Postdoctoral Fellow program (Marie Skłodowska-Curie Co. funded, Grant 754462). Serge Söntjes is kindly acknowledged for assistance with the initial NMR experiments. Xavier Guillory is also thanked for the inspiration for this manuscript.

