## Supporting information for "Early identification of cooperative fragments for protein-protein interaction stabilization"

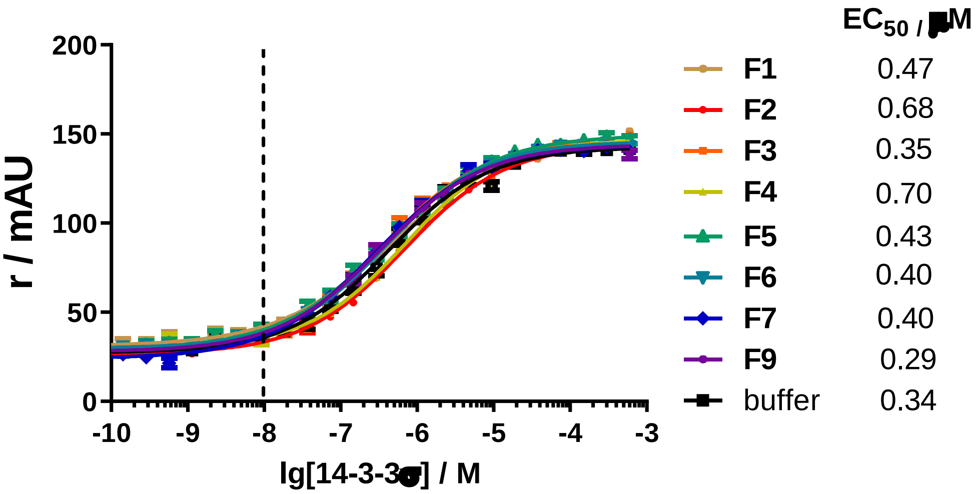


**Supporting Figure S1.** FA measurements of 14-3-3σ to 10 nM FITC-labeled TAZ peptide in the presence of 1mM fragments in FA-buffer (10 mM HEPES, 150 mM NaCl, 1 mg/ml BSA 0.01%, v/v TWEEN-20, and 2% DMSO). Error bars indicate standard deviation (n=3).

**Supporting Table S1.** Signal intensity

| Fragment | Peak (ppm) | Signal intensity | |
|---|---|---|---|
| | | without peptide (SNR) [b] | with peptide (SNR) |
| **AZ130** | 7,69 (s) | 2.91 | 6.09 |
| **F1** | 7,15 (s) | 6.39 | 2.83 |
| **F2** | 8,72 (s) | 1.55 | 8.93 |
| **F3** | NS [a] | NS | NS |
| **F4** | 7,16 (s) | 3.04 | 5.45 |
| **F5** | 8,32 (s) | 4.94 | 11.31 |
| **F6** | NS | NS | NS |
| **F7** | NS | NS | NS |
| **F8** | 6,96 (d) | 5.15 | 4.03 |
| **F9** | 7,86 (s) | NS | 3.67 |

[a] NS indicates no signal was detected. b SNR; signal-to-noise-ratio. Value was calculated based on the peak intensity of an aromatic singlet relative to the baseline of the STD NMR base line. One exception was **F8** were a doublet was selected.

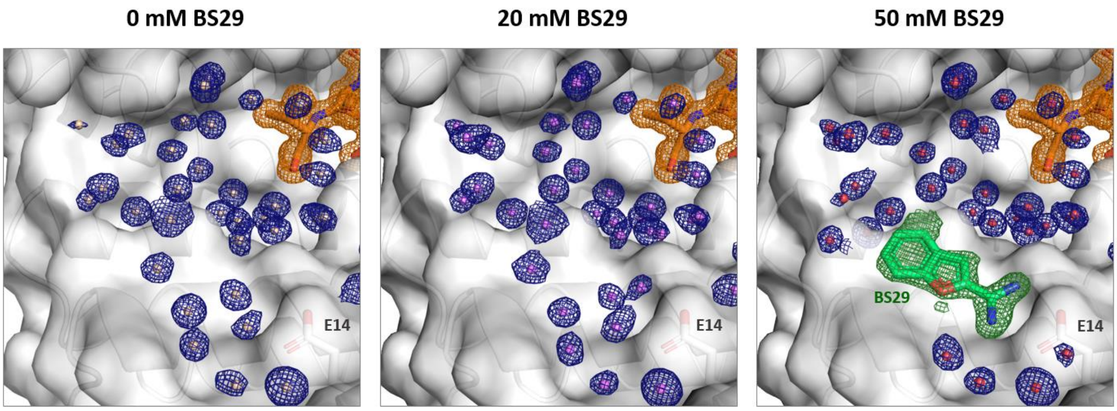


**Supporting Figure S2.** Crystal structure of ERα (orange sticks and electron density) in complex with 14-3-3σ (grey surface) soaked with different concentrations of **F9** (=BS29) (green sticks and electron density). Water molecules are illustrated as small spheres with their correlated electron density maps. All $2F_0$-$F_c$ electron density maps are contoured at 1σ.

**Supporting Equation1.** Amplification factor ($A_{STD}$) equation.

(1) $$A_{STD} = \frac{I_0 - I_{SAT}}{I_0} \times \frac{[L]_T}{[P]} = \frac{I_{STD}}{I_0} \times \frac{[L]_T}{[P]}$$

Where;

$I_0$ = equilibrium intensity (off-resonance)
$I_0$ = saturation intensity (on-resonance)
$[L]_T$ = Ligand concentration at a set saturation time
[P] = Protein concentration

#### Materials and Methods

##### Compounds

All fragments were bought from enamine and structures confirmed using LCMS, $^{1}H$ and $^{13}C$ NMR.

##### Peptides

All peptides were procured via Genscript and used as is.

| Name | Binding Site | Sequence |
|---|---|---|
| TAZ | pS89 | *Ac*-RSH pS SPASLQLGT-*CONH₂* |
| TASK3 | pS373 | *Ac*-KRRKpSV-*COOH* |
| ABL1 | pT735 | *Ac*-EWRSV pT LPRDL-*CONH₂* |
| AS160 | pT642 | *Ac*-RRRAH pT FSHPP-*CONH₂* |
| TBC1D | pS237 | *Ac*-MRKSF pS QPGLR-*CONH₂* |
| Synaptopodin | pS137 | *Ac*-LRLAY pS EPCGL-*CONH₂* |
| REDD1 | pS2126 | *Ac*-PSRAK pS RPLPN-*CONH₂* |
| FITC-TAZ | pS89 | FITC-O1Pen-RSH pS SPASLQLGT-*CONH₂* |

##### Protein expression and purification used for NMR experiments

A pPROEX HTb expression vector encoding the human 14-3-3 protein sigma (14-3-3σ) with an N-terminal $His_6$-tag was transformed by heat shock into NiCo21 (DE3) competent cells. Site-directed mutagenesis to obtain E14A mutant was performed using the QuickChangeLightning site-direct mutagenesis kit (Agilent Technologies) following manufacturer's instructions (Forward primer: GAA GGC CAA GCT GGC AGC ACA GGC CGA ACG CTA TG; Reverse primer: CAT AGC GTT CGG CCT GTG CTG CCA GCT TGG CCT TC). Successful mutagenesis was confirmed by DNA sequencing.

Single colonies (wt or mutated 14-3-3) were cultured in 50 mL LB medium (10 μg/mL ampicillin). After overnight incubation at 37 °C, cultures were transferred to 2 L TB media (10 μg/mL ampicillin, 1 mM $MgCl_2$) and incubated at 37 °C until an OD600 nm of 0.8-1.2 was reached. Protein expression was then induced with 0.4 mM isopropyl-β-d-thiogalactoside (IPTG), and cultures were incubated overnight at 18˚C. Cells were harvested by centrifugation (8600 rpm, 20 minutes, 4 ˚C) and resuspended in lysis buffer (50 mM HEPES, pH 8.0, 300 mM NaCl, 12.5 mM imidazole, 5 mM $MgCl_2$, 2 mM βME) containing cOmplete™ EDTA-free Protease Inhibitor Cocktail tablets (1 tablet/ 100 ml lysate) and benzonase (1 µl/ 100 ml). After lysis using a C3 Emulsiflex-C3 homogenizer (Avestin), the cell lysate was cleared by centrifugation (20000 rpm, 30 minutes, 4 ˚C) and purified using $Ni^{2+}$-affinity chromatography (Ni-NTA superflow cartridges, Qiagen). Typically two 5 mL columns (flow 5 mL/min) were used for a 2 L culture in which the lysate was loaded on the column washed with 10 CV wash buffer (50 mM HEPES, pH 8.0, 300 mM NaCl, 25 mM imidazole, 2 mM βME) and eluted with several fractions (2-4 CV) of elution buffer (50 mM HEPES, pH 8.0, 300 mM NaCl, 250 mM imidazole, 2 mM βME). Fractions containing the 14-3-3 protein were combined and dialyzed into 25 mM HEPES pH 8.0, 100 mM NaCl, 10 mM $MgCl_2$, 500 µM TCEP. Finally, the protein was concentrated to ~60 mg/mL, analyzed for purity by SDS-PAGE and Q-Tof LC/MS and aliquots flash-frozen for storage at −80 °C.

##### Protein expression and purification used for crystallography

A pPROEX HTb expression vector encoding the human 14-3-3 protein sigma truncated after T231 (14-3-3σ Δc) and with an N-terminal $His_6$-tag was transformed by heat shock into NiCo21 (DE3) competent cells. Single colonies were cultured in 50 mL LB medium (10 μg/mL ampicillin). After overnight incubation at 37 °C, cultures were transferred to 2 L TB media (10 μg/mL ampicillin, 1 mM $MgCl_2$) and incubated at 37 °C until an OD600 nm of 0.8-1.2 was reached. Protein expression was then induced with 0.4 mM isopropyl-β-d-thiogalactoside (IPTG), and cultures were incubated overnight at 18˚C. Cells were harvested by centrifugation (8600 rpm, 20 minutes, 4 ˚C) and resuspended in lysis buffer (50 mM HEPES, pH 8.0, 300 mM NaCl, 12.5 mM imidazole, 5 mM $MgCl_2$, 2 mM βME) containing cOmplete™ EDTA-free Protease Inhibitor Cocktail tablets (1 tablet/ 100 ml lysate) and benzonase (1 µl/ 100 ml). After lysis using a C3 Emulsiflex-C3 homogenizer (Avestin), the cell lysate was cleared by centrifugation (20000 rpm, 30 minutes, 4 ˚C) and purified using $Ni^{2+}$-affinity chromatography (Ni-NTA superflow cartridges, Qiagen). Typically two 5 mL columns were used for a 2 L culture in which the lysate was loaded on the column washed with 10 CV wash buffer (50 mM HEPES, pH 8.0, 300 mM NaCl, 25 mM imidazole, 2 mM βME) and eluted with several fractions (2-4 CV) of elution buffer (50 mM HEPES, pH 8.0, 300 mM NaCl, 250 mM imidazole, 2 mM βME). Fractions containing the 14-3-3 protein were combined and dialyzed into 25 mM HEPES pH 8.0, 200 mM NaCl, 10 mM $MgCl_2$, 2 mM βME. In addition, 1 mg TEV was added for each 100 mg purified protein to remove the purification tag. The cleaved sample was then again loaded on a 10 mL Ni-NTA column to separate the cleaved product from the expression tag and residual uncleaved protein. The flowthrough was loaded on a Superdex 75 pg 16/60 size exclusion column (GE Life Sciences) using 25 mM HEPES, 100 mM NaCl, 10 mM $MgCl_2$, 500 µM TCEP (adjusted to pH=8.0) as running buffer. Fractions containing the 14-3-3 protein were pooled and concentrated to ~60 mg/mL, analyzed for purity by SDS-PAGE and Q-Tof LC/MS and aliquots flash-frozen for storage at −80 °C.

14-3-3σ full length (1-248)
SYYHHHHHHDYDIPTTENLYFQGAMGSMERASLIQKAKLAEQAERYEDMAAFMKGAVEKGEELSCEERNLLSVAY
KNVVGGQRAAWRVLSSIEQKSNEEGSEEKGPEVREYREKVETELQGVCDTVLGLLDSHLIKEAGDAESRVFYLKMK
GDYYRYLAEVATGDDKKRIIDSARSAYQEAMDISKKEMPPTNPIRLGLALNFSVFHYEIANSPEEAISLAKTTFDEAM
ADLHTLSEDSYKDSTLIMQLLRDNLTLWTADNAGEEGGEAPQEPQS

14-3-3σ full length E14A (1-248)
SYYHHHHHHDYDIPTTENLYFQGAMGSMERASLIQKAKLAAQAERYEDMAAFMKGAVEKGEELSCEERNLLSVAY
KNVVGGQRAAWRVLSSIEQKSNEEGSEEKGPEVREYREKVETELQGVCDTVLGLLDSHLIKEAGDAESRVFYLKMK
GDYYRYLAEVATGDDKKRIIDSARSAYQEAMDISKKEMPPTNPIRLGLALNFSVFHYEIANSPEEAISLAKTTFDEAM
ADLHTLSEDSYKDSTLIMQLLRDNLTLWTADNAGEEGGEAPQEPQS

14-3-3σ ΔC (1-231)
SYYHHHHHHDYDIPTTENLYFQ|GAMGSMERASLIQKAKLAEQAERYEDMAAFMKGAVEKGEELSCEERNLLSVA
YKNVVGGQRAAWRVLSSIEQKSNEEGSEEKGPEVREYREKVETELQGVCDTVLGLLDSHLIKEAGDAESRVFYLKMK
GDYYRYLAEVATGDDKKRIIDSARSAYQEAMDISKKEMPPTNPIRLGLALNFSVFHYEIANSPEEAISLAKTTFDEAM
ADLHTLSEDSYKDSTLIMQLLRDNLTLWT
C-terminal truncation is made to improve crystallization of 14-3-3.

….. = purification tag
| = TEV cleavage site
A = mutated site

**STD-NMR**

General protocol

NMR samples for STD experiments were prepared with 10 µM 14-3-3 protein, 100 µM peptide and 300 µM fragment, in PBS (50 mM $NaH_2PO_4/Na_2HPO_4$, 150 mM NaCl, pH 7.5) with 10% $D_2O$ (v/v). Standard 1D and STD NMR spectra were acquired at 20 °C with a 400 MHz Bruker Avance 400 MHz NMR.

The STD spectra were measured by using a 1D STD with spoil and T2 filter using excitation sculpting (stddiffesgp.3) pulse program. A shaped pulse train for saturation on f2 channel was used alternating between on- and off resonance. The saturation time (D20) was set to 2 s and the spin lock time of the protein background (D29) set to 25 ms. The relaxation delay (D1) was set to equal the saturation time. The on-resonance frequency used for saturation was set to 2.0 ppm, while the off-resonance irradiation was applied at 16 ppm, where no NMR resonances were present. Dummy scans (DS) and number of scans (NS) were set to 4 and 1024, respectively. The STD spectra were processed with Bruker Topspin 3.5 pl 5.

Initial fragment screen
10 $\mu$M 14-3-3σΔC with 100 $\mu$M peptide and 300 $\mu$M fragment.

STD signal intensity was determined using Singal Noise Ratio (SNR) peak calculator script in MestReNova© 11.0.4-18998 software package. This script calculates the ration of signal relative to the baseline. For all fragments, with the exception **F8**, a singlet signal was selected for integration and the region -1 to -3 ppm was selected as the baseline region. This procedure was conducted for both with and without peptide.

Validation
10 $\mu$M 14-3-3σΔC or 10 $\mu$M 14-3-3σΔC_E14A with 100 $\mu$M peptide and 5 mM fragment

Amplification STD ($A_{STD}$) values were processed using Bruker TopSpin 3.5pl5 software package and the experimental procedure described in the supporting information of *Viegas et al.*[1]

Selectivity screen
10 $\mu$M 14-3-3σΔC with 100 $\mu$M peptide and 5mM fragment

Amplification STD ($A_{STD}$) values were processed using Bruker TopSpin 3.5pl5 software package and the experimental procedure described in the supporting information of *Viegas et al.*[1]

**Fluorescence anisotropy measurements**
Fluorescence anisotropy affinity measurements were conducted in FA-buffer (10 mM HEPES pH 7.4, 150 mM NaCl) with 0.01% TWEEN-20 and 1.0 mg/mL BSA, using fixed concentrations of fluorescently-labeled peptide (10 nM), a fixed concentration of 1 mM fragment and DMSO (2% v/v) in 10 µL round-bottom low binding 384-micro well plates (Corning, #25916024). Measurements were performed in triplicate at 20 °C with a filter-based microplate reader (Tecan Infinite F500) using a fluorescein filterset (λex: 485 nm/20 nm, λem: 535 nm/25 nm) and an integration time of 50 µs, errors bars indicate the standard deviations. The data was fitted using GraphPad Prism 5.05 for Windows (GraphPad Software Inc., CA, USA four-parameter logistic model (4PL) to obtain the $EC_{50}$-values.

**X-ray crystallography data collection and refinement**
ERα-8mer (Ac-AEGFPApTV-COOH) peptide and 14-3-3σΔC protein were dissolved in complexation buffer (20 mM HEPES pH 7.5, 100 mM NaCl, 10 mM $MgCl_2$ and 20 µM TCEP) using a final 14-3-3 concentration of 12.5 mg/mL and a 1:2 molar stoichiometry of protein:peptide. These complexes were incubated overnight at room temperature. After the incubation, sitting-drop crystallization plates were set up in which each of the four complexation mixtures was combined with 24 crystallization buffers, optimized for 14-3-3σ crystallization (0.095 M HEPES pH 7.3, 0.19 M $CaCl_2$, 26 (v/v) PEG 400 and 5% (v/v) glycerol). Herein a 1:1 mix (both 250 nL) of complexation mixture and crystallization buffer was made for crystal growth. Crystals grew within 10 - 14 days at 4 °C.

Soaking crystals was performed by mixing 2.0 μL of 100 mM fragment stock solutions in DMSO with 2.0 μL mother liquor, and adding this to crystal-containing drops. Crystals were soaked for 6 days after they were fished and flash-cooled in liquid nitrogen. X-ray diffraction (XRD) data were collected either at the Deutsches Elektronen-Synchortron (DESY, PETRA-III, beamline P11, Hamburg, Germany) equipped with a PILATUS 6M-F detector. Typical settings were 1440 image, 0.25°/image, 10% transmission and 0.1 s exposure time.

Data was processed using the CCP4i2 suite (version 7.1.10).[2] DIALS[3] was used to index and integrate the data after which scaling was done using AIMLESS.[4,5] The data was phased with MolRep[6], using protein data bank (PDB) entry 4JC3 as a template. Ligand restraints for non-natural amino acids were generated with eLBOW[7]. Sequential model building (based on visual inspection Fo-Fc and 2Fo-Fc electron density map) and refinement were performed with COOT and REFMAC, respectively.[8-10] Finally, alternating cycles of model improvement and refinements were performed using coot and phenix.refine from the Phenix software suite (version 1.15.2-3472).[11,12] Pymol (version 2.2.3)[13] was used to make the figures and the structures were deposited in the protein data bank (PDB) with ID: 7O2J (See Table S1 for crystal statistics).

**Table S1.** Data collection and refinement statistics (molecular replacement) for 14-3-3σΔc in complex with Estrogen Receptor α phosphopeptide and fragment **9** (PDB: 7O2J)

| | 14-3-3σΔc/ERα/Fragment 9 |
|---|---|
| *Data collection* | |
| Space group | C 2 2 21 |
| Cell dimensions | |
| a, b, c (Å) | 82.21, 112.59, 62.60 |
| α, β, γ (°) | 90, 90, 90 |
| Resolution (Å) | 45.54 – 1.50 (1.54 – 1.50) |
| $I / \sigma(I)$ | 22.8 (6.6) |
| Completeness (%) | 100.0 (100.0) |
| Redundancy | 12.7 (12.7) |
| $CC_{1/2}$ | 0.99 (0.971) |
| *Refinement* | |
| No. reflections | 46809 |
| $R_{work}/R_{free}$ | 0.157/0.173 |
| No. atoms | |
| Protein | 1949 |
| Ligand/ion | 15 |
| Water | 358 |
| *B*-factors | |
| Protein | 13.79 |
| Ligand/ion | 27.82 |
| Water | 26.41 |
| R.m.s. deviations | |
| Bond lengths (Å) | 0.008 |
| Bond angles (°) | 0.96 |

### STD NMR Spectra with TAZ binding peptide

#### Fragement 1 (F1)

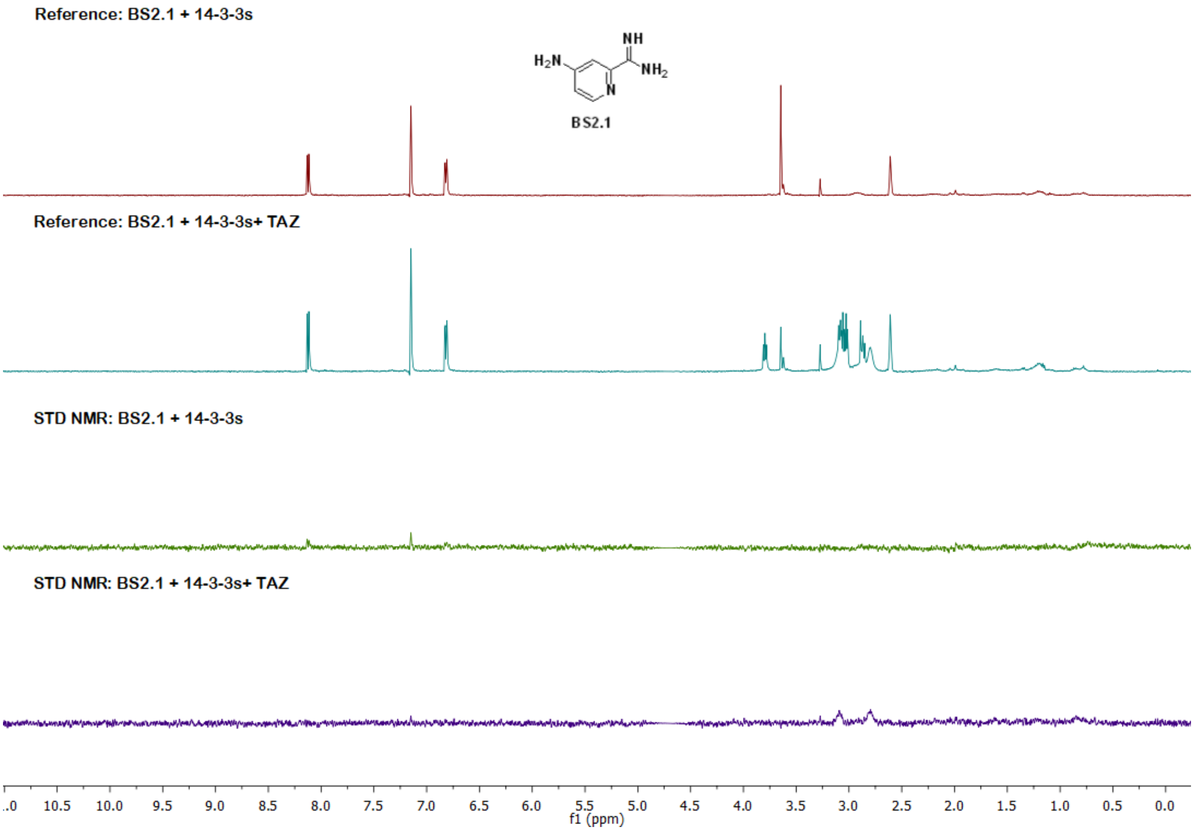


#### Fragement 2 (F2)

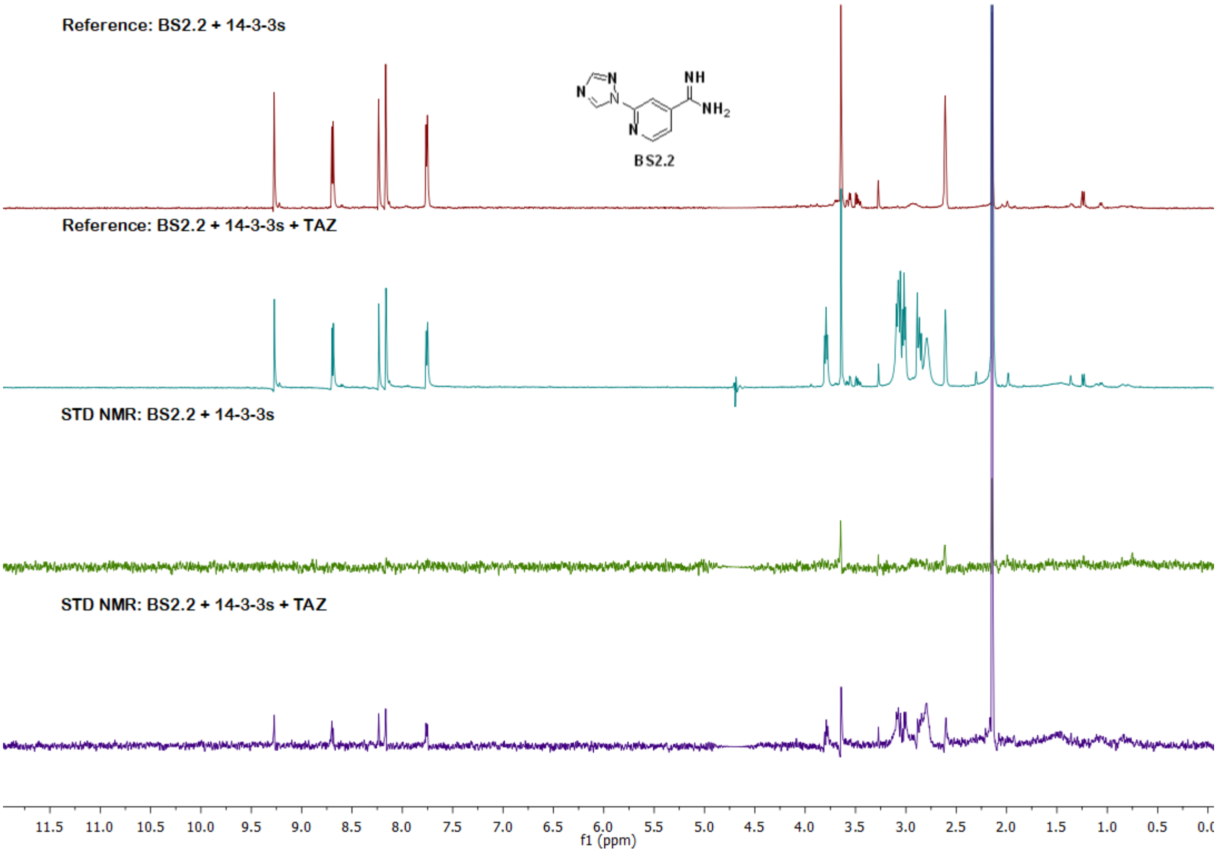

#### Fragement 4 (F4)

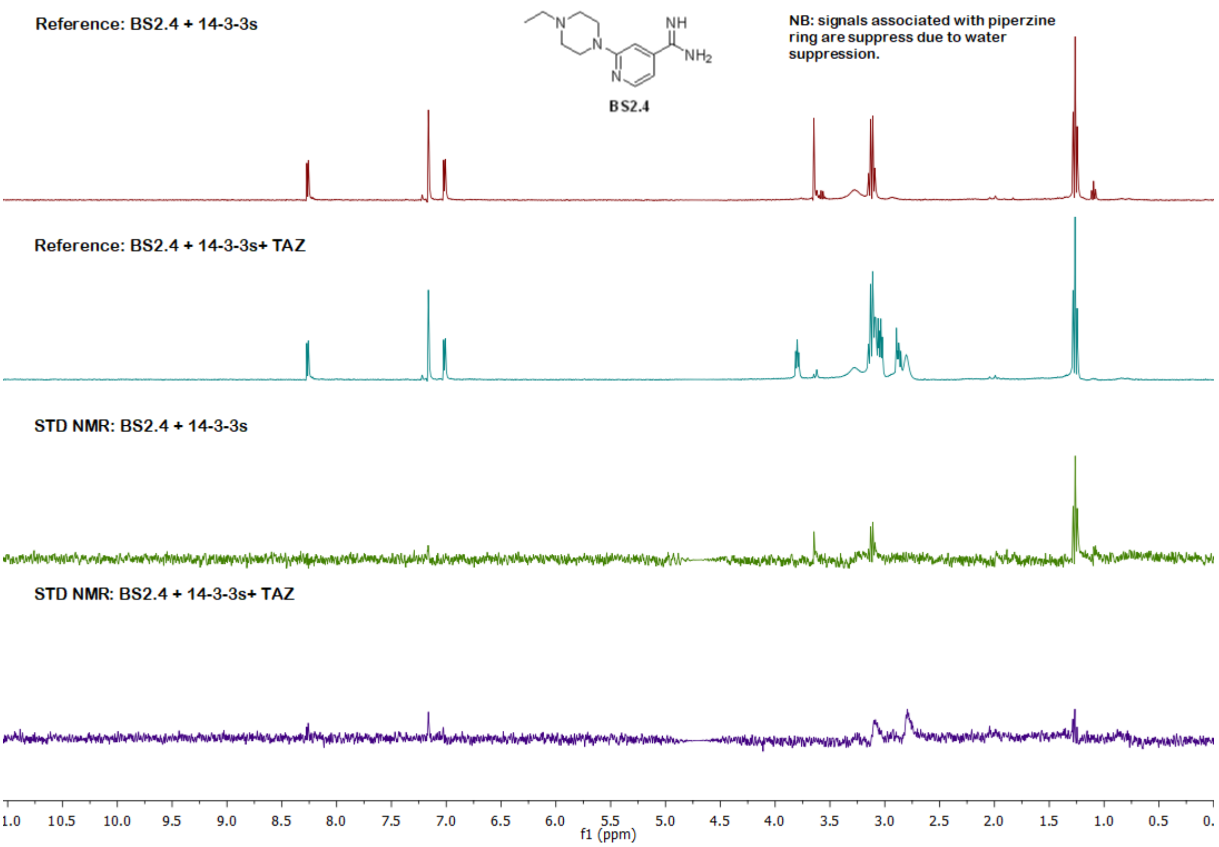


#### Fragement 5 (F5)

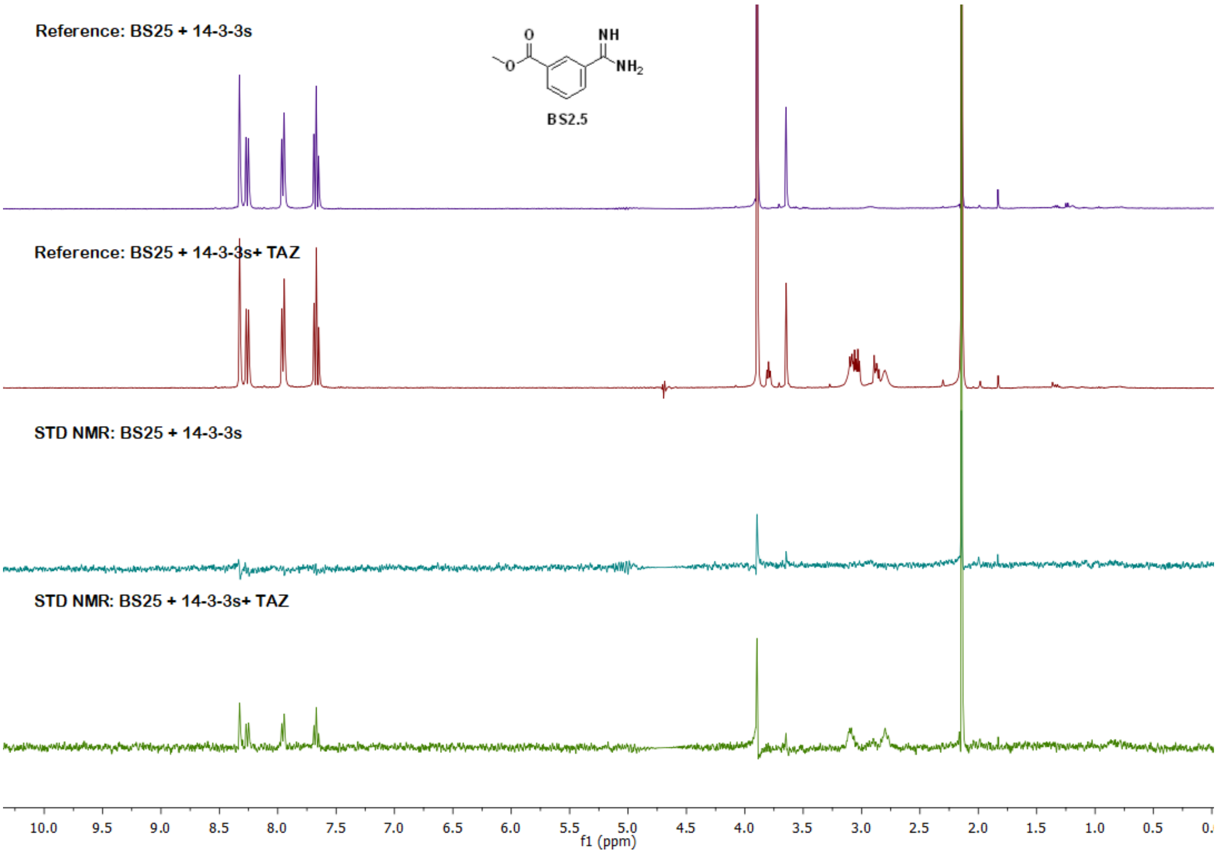

#### Fragement 6 (F6)

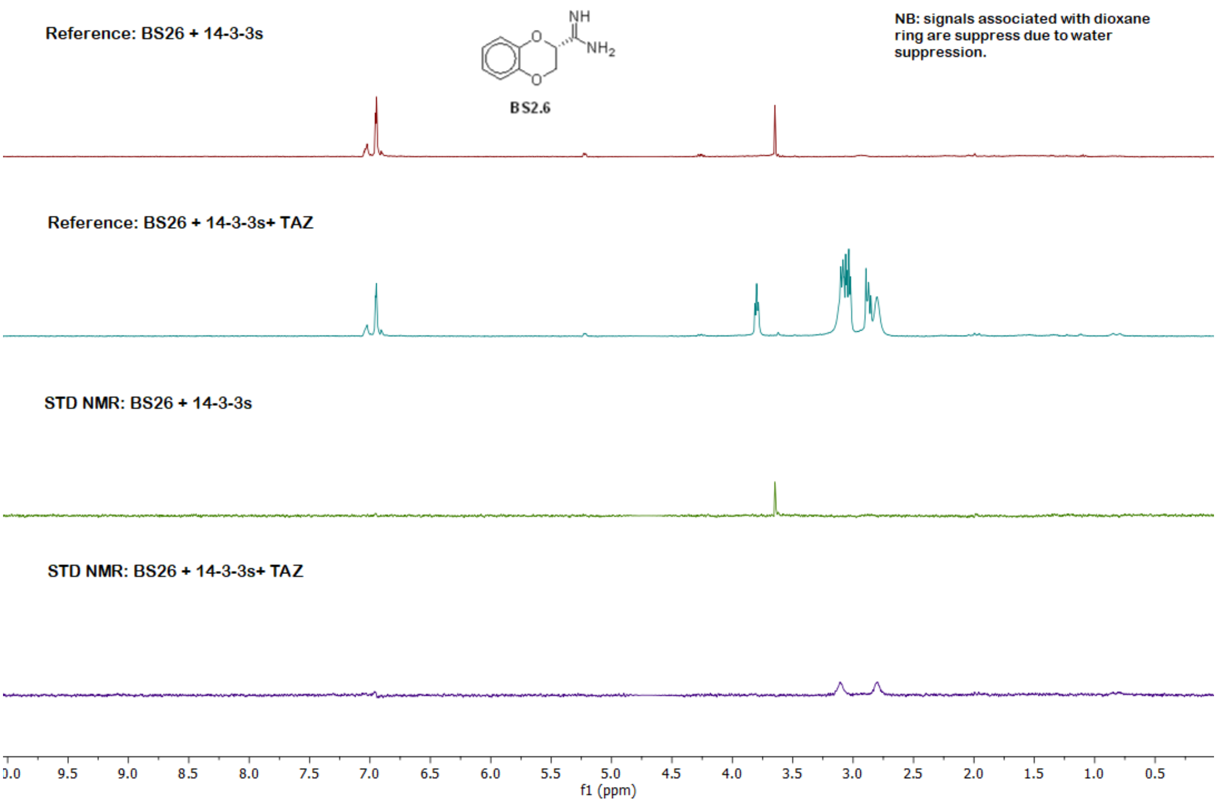


#### Fragment 7 (F7)

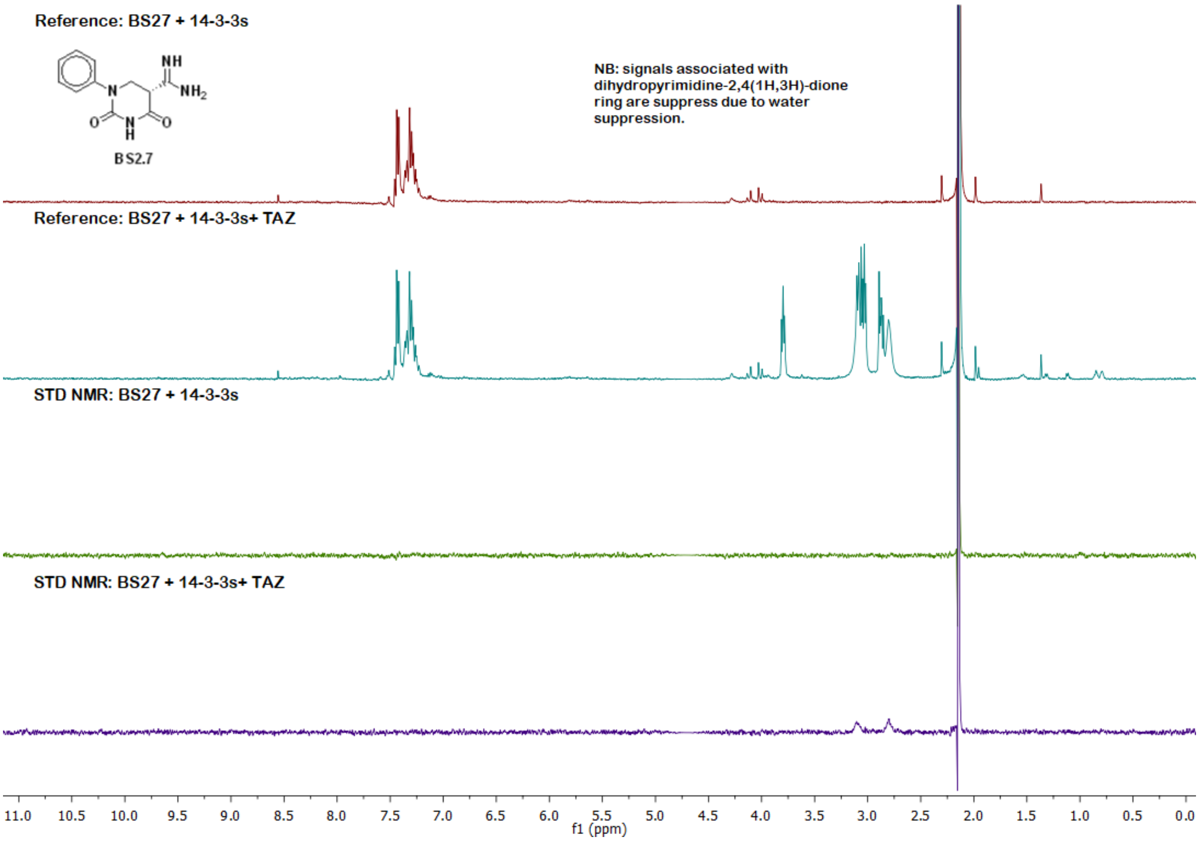

#### Fragment 8 (F8)

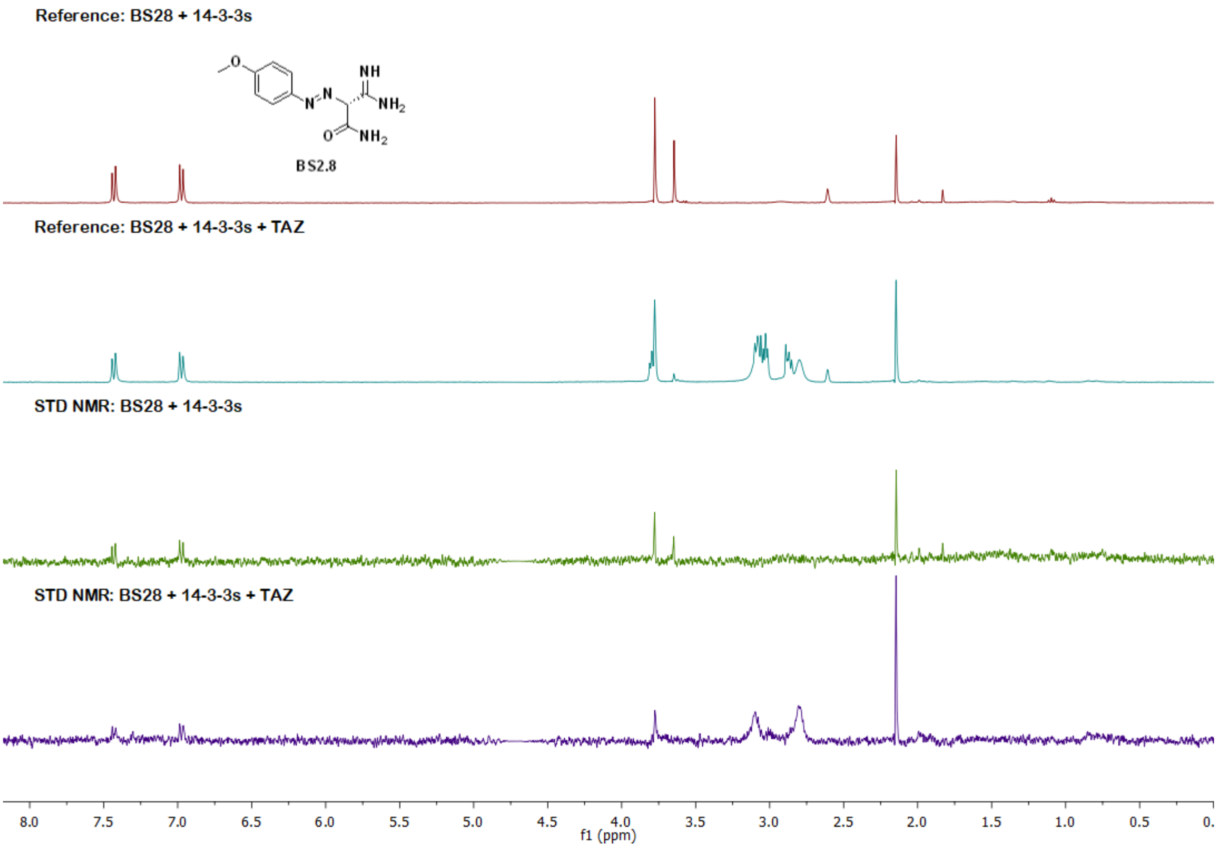


#### Fragment 9 (F9)

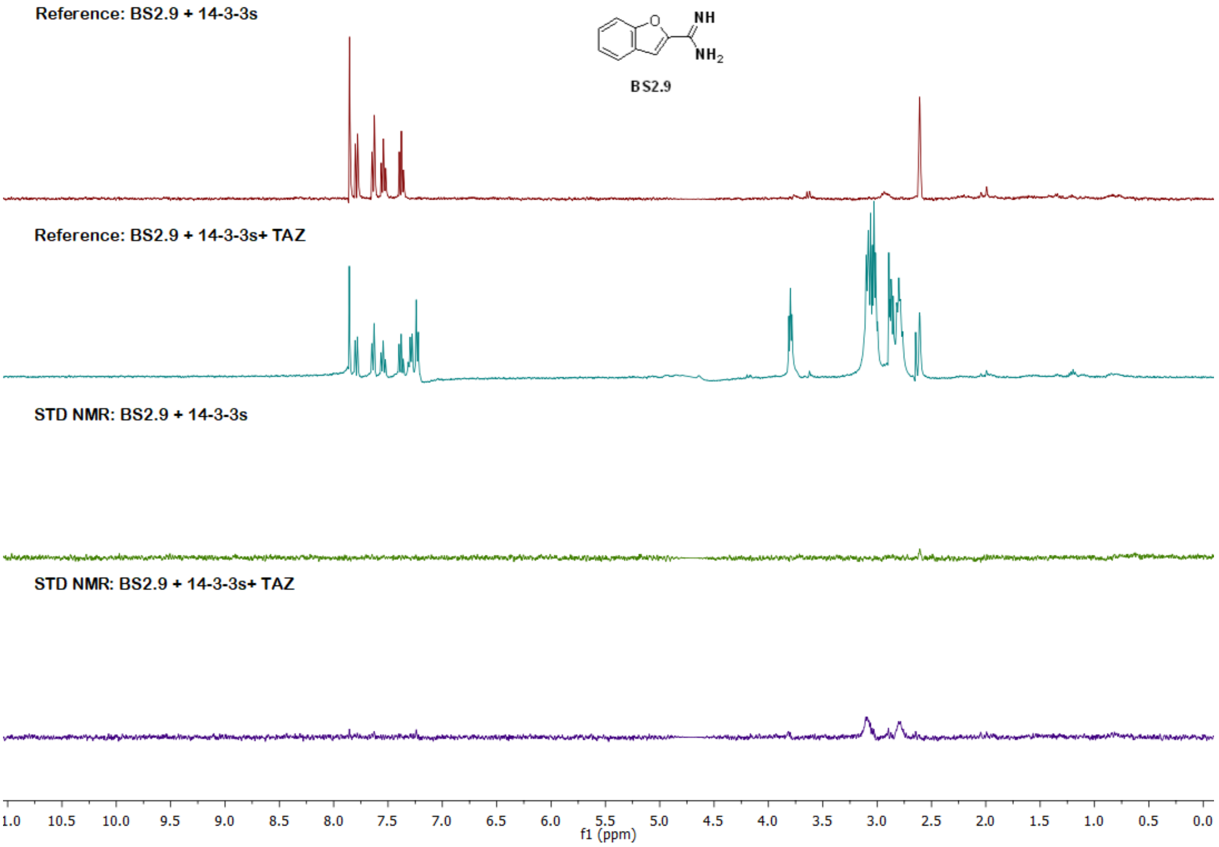
